# Genes associated with liver damage signalling pathways may impact the severity of COVID-19 symptoms in Spanish and Italian populations

**DOI:** 10.1101/2020.07.03.179028

**Authors:** Leire Moya, Samaneh Farashi, Prashanth N Suravajhala, Panchadsaram Janaththani, Jyotsna Batra

**Author notes:** Correspondence should be addressed to: A/Prof Jyotsna Batra, Australian Prostate Cancer Centre-Queensland, Translational Research Institute, Queensland University of Technology, Woolloongabba, Queensland, Australia 4102.

## Abstract

**Aim:** The novel SARS-CoV-2 virus, which causes the COVID-19 disease, has infected more than 10 million people and caused 500K deaths worldwide. In Europe, over 2 million confirmed cases have been reported, while nearly 200K people have died from the disease. Despite strict containment measures in Spain and Italy after the first reported COVID-19 patient, these two countries have remained in the top five European nations with the highest mortality rate for over two months. We hypothesised that a genetic mechanism could partially explain the poor survival outcome observed in these two countries.

**Methods:** An extensive literature search to identify human candidate genes linked to SARS-CoV infection, host immune evasion and disease aggressiveness was carried out. Pathway analysis (IPA) was performed to select the most significantly associated canonical signalling pathways with the genes of interest. The genetic variants’ at these genes with ±1Mb flanking region was extracted (GRCh37/hg19 built). Over 80 million single nucleotide polymorphisms (SNPs) were analysed in genome-wide data of 2,504 individuals (1000 genomes, phase III, https://www.internationalgenome.org/). Principal component (PC) analysis was performed, ancestry by the whole genome was inferred and subsets of the regions of interest were extracted (PLINK v1.9b, http://pngu.mgh.harvard.edu/purcell/plink/). PC1 to PC20 values from five European ancestries, including the Spanish and Italian populations, were used for PC analysis. Gene function predictions were run with our genes of interest as a query to the GeneMANIA Cytoscape plugin (https://genemania.org/).

**Results:** A total of 437 candidate genes associated with SARS were identified, including 21 correlated with COVID-19 aggressiveness. The two most significant pathways associated with all 437 genes (*Caveolar-mediated Endocytosis* and *MSP-RON Signalling*) did not show any segregation at the population level. However, the most significant canonical pathway associated with genes linked to COVID-19 aggressiveness, the *Hepatic Fibrosis and Hepatic Stellate Cell Activation,* showed population-specific segregation. Both the Spanish and Italian populations clustered together from the rest of Europe. This was also observed for the Finnish population but in the opposite direction. These results suggest some of the severe COVID-19 cases reported in Spain and Italy could be partially explained by a pre-existing liver condition (especially liver cancer) and/or may lead to further COVID-19 related liver complications.

## Introduction

The novel coronavirus disease 19 (COVID-19), is an infectious disease caused by the severe acute respiratory syndrome coronavirus 2 (SARS-CoV-2). It was firstly identified in December 2019 [1] and since then, nearly 500K people have died worldwide global deaths (27^th^ June 2020 [2]). SARS-CoV-2 is closely related to SARS-CoV-1, which was firstly identified in November 2002 in China [3]. Despite the virus spreading to 26 countries [4], only five localised hotspots worldwide remained [5], and by July 2003 the virus outbreak was declared officially contained by the World Health Organisation (WHO) [6]. A total of 774 people died [3] and no new cases have been reported since 2004 [7]. On that year, R_0_ data from the three main hotspots (Singapore, Toronto and Hong Kong) showed SARS-CoV-1 had an R_0_ = 0.9 – 1.8 [8]. The R_0_ value is and indicative of how an epidemic can spread in a completely susceptible population where no specific control measures are implemented. The higher the value, the more difficult is for the population to contain the spread. Data from February and April this year, report a mean value of R_0_=2·6 [9, 10] for SARS-CoV-2. This data explains the broad success of the containment measures implemented during the first SARS pandemic [4], where the localisation of the virus lead to its eradication.

At the end of February 2020 (29^th^), Italy recorded the highest number of community transmitted COVID-19 related deaths (21 total deaths [11]) in Europe. Two weeks later, Spain was the second European county with the highest COVID-19 related deaths (136 deaths, [11]) and less than a month later, both countries had 29.3 and 28.3 deaths per 100,000 respectively, while the rest of Europe reported 0.45 – 11.3 deaths (7^th^ of April, [11]). Currently (4^th^ of June 2020), Belgium and the UK have reported the highest COVID-19 mortality rate while Spain and Italy remain the third and fourth countries [11]. The reported COVID-19 mortality rates are most likely the consequences of a combination of factors such as the Public Health’s response, social behaviour and population comorbidities.

As such, we searched what the differences in terms of the responses to the crisis were for these four countries. Both in Spain and Italy, strong containment measures were implemented four and three weeks after the first confirmed case in their countries [12]. Belgium started its most strict rules five weeks after the first case while it took six weeks for the UK to take action [12]. Similarly, several COVID-19 comorbidities have been identified such as aging, diabetes, heart and liver diseases [13–16]. But little is known about which genes and/or pathways are associated with the increased risk of developing severe symptoms.

At the peak of the pandemic in Italy and Spain, we hypothesised that a genetic component in these populations may contribute to the high mortality rate of COVID-19 patients observed. To study this, we performed pathway analysis of 437 candidate genes associated with SARS-like viral intake, immune evasion and clinical phenotype, which were shortlisted from the extensive literature study. Principal Component (PC) analysis of the single nucleotide polymorphisms (SNPs) and their allele frequencies in these candidate gene loci with a 1Mb window was performed. Our analysis revealed both Italian and Spanish populations segregate differently in the pathways involved in liver injury and damage, with a potential role of the hepatic disease being involved in a more aggressive response to the COVID-9 disease.

## Material and methods

### Mortality rate frequency calculation

The *cause-specific death rate* was measure using the below formula provided by the Centers for Disease Control and Prevention [17] and the data is presented per 100,000 individuals. The total *COVID-19 associated deaths* were defined by the International Guidelines defined by WHO (based on ICD) [18] and the population size was sourced from the Eurostat Data Browser [19] as of 1^st^ of January 2020.

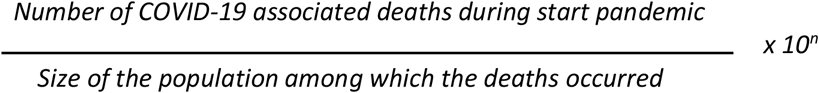

Public health data from the European populations studied here was searched regarding the most common COVID-19 comorbidities reported. Aging has been broadly identified as the main one [13, 20, 21], followed by cardiovascular disease (CVD), diabetes, chronic obstructive pulmonary disease (COPD) and cancer history [13, 15]. Liver dysfunction, smoking status, chronic kidney diseases and immunodeficiency have also been reported [14–16].

### Identification of candidate genes for analysis

PubMed was accessed between the 31^st^ of March and 25^th^ of May 2020. The aim was to consolidate all empirically and predicted human genes reported to have a role during SARS-CoV-2 infection to date. This included those human genes that assisted with the viral entry, evasion of the host’s immune system and the SARS-CoV-2 − human interactome (Figure 1). Those studies which results did not include human genes interacting direct or indirectly with SARS-CoV-1 or SARS-CoV-2 or being affected by the disease they cause were excluded.

**Figure 1:**
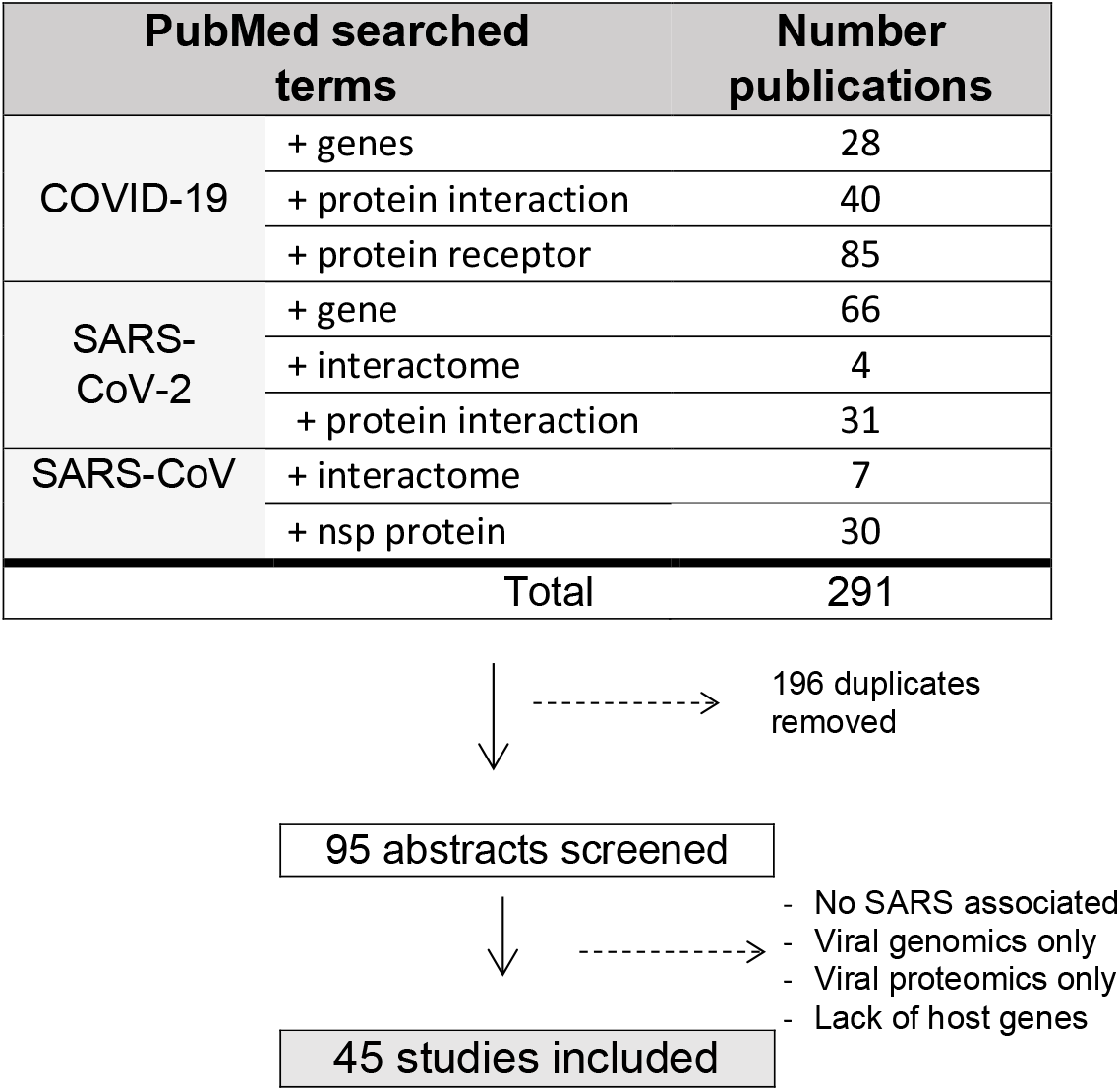
PubMed search between the 31st of March and 25th of May 2020. Several searching terms were entered in the database to identify all reported SARS-CoV host interactome reported genes.

**Figure 2:**
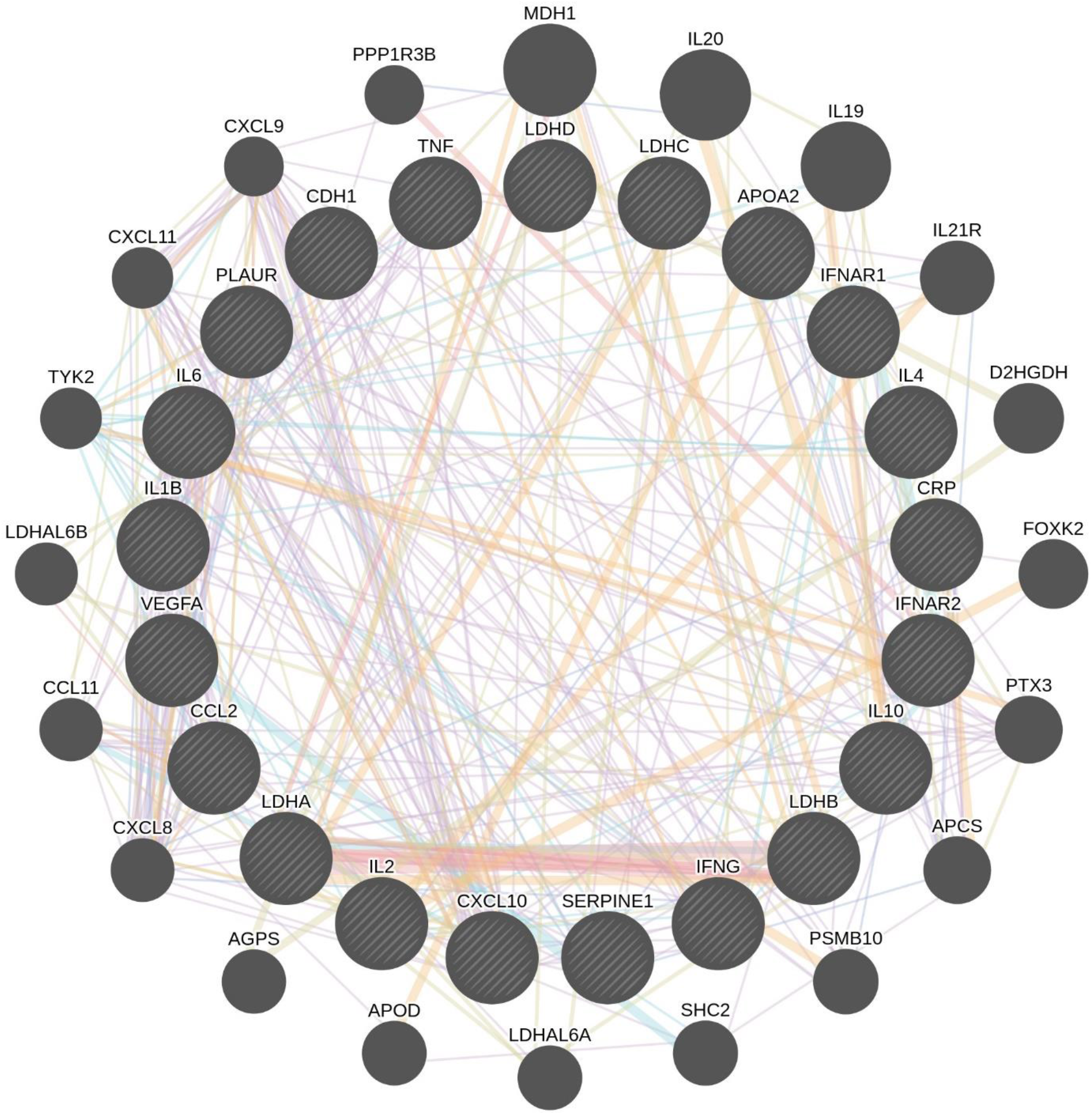
Protein-Protein Interaction map of all the 21 candidate genes associated with COVID-19 aggressiveness from our pilot study. The pin edge syndicate the amount of physical interactions that are well known from public literature with LDHA and LDHB as key players. Analysis run with GeneMANIA (https://genemania.org/). LDH: lactate dehydrogenase

After completing the literature research, all the genes identified were curated for the subsequent pathway and PC analyses. For example, when the literature reference did not specify the gene isoform (e.g. NF-κB) all those available (e.g. NFKB1 and NFKB2) where included in the analysis. Pathway signalling analysis was performed with IPA software (QIAGEN Inc., [22]) as follows. All candidate genes’ symbols, or subtracted genes as described below, were entered for the Core Analysis Expression. As a reference, the *Ingenuity Knowledge Base* was applied and both *Direct and Indirect Analysis Analyses* were selected. The genes involved in the two most significant canonical pathways affected by our candidate genes’ list input were selected and their locations in the genome were identified after accessing the *Table Browser* tool from UCSC (https://genome.ucsc.edu/index.html) (GRCh37/hg19 built). Duplicates were removed while unique transcript variants for a gene were included. The flanking regions of ± 1Mb were added wand the allele frequencies in these loci were analysed for further PC analysis amongst the populations of interest.

Also, selected genes associated with COVID-19 severe symptoms were entered in the GeneMANIA Cytoscape plugin to predict physical protein-protein interactions [23]. This plugin includes a large dataset of over 800 functional associated networks for human amongst other organisms. The data utilised come from reported studies and public large databases such as BIOGRID (Breitkreutz et al., 2008), GEO (Barrett et al., 2009), I2D (Brown and Jurisica, 2005) and Pathway Commons (http://www.pathwaycommons.org). The regular and automated update of the networks makes this tool and up to date resource for the category analysed.

### PC Analysis

The genome-wide germline data of 2,506 individuals were obtained from 1000 genomes phase III v.5b project, the latest release of the data (May 2013) (https://www.internationalgenome.org/). Starting from the genotype file vcf.gz formats provided by 1000 genomes projects, we performed standard quality control methods using PLINK v1.9b (http://pngu.mgh.harvard.edu/purcell/plink/) [24] by removing individuals with more than 3% missing genotypes, SNPs with a call rate <97%.

We performed PC analysis for the samples to account and adjust for population stratification. PCs were computed using PLINK software v1.9b for all samples of 1000G (1KG) project, phase 3 version 5b, to infer the ancestry of the samples based on the whole genome data. We used PLINK [24] to extract subsets of regions for 1Mb window of genes of our interest. PC1 and PC20 values >6 standard deviations from five European ancestries: Great Britain (England and Scotland), CEU (Utah Residents with Northern and Western European Ancestry), Tuscany (Italy), Iberian (Spanish population) and Finland participants were used in the analysis.

## Results

### Mortality rate and common COVID-19 comorbidities data in European populations

Calculations of the mortality rates reported in Europe were done for the 27 countries that form the European Union [25] as shown in the Methods section. This was first calculated at the beginning of the study (7^th^ of April) and updated later to reflect any changes (4^th^ of June Table 1). Despite the recent separation of the UK from the EU, we included it for its relevancy towards this study. Since April, Spain, Italy and Belgium have remained in the top five countries with the highest mortality rate (Table 1).

**Table 1:** COVID-19 mortality rate per 100,000 people in the European Union.

|  | Country | Deaths in<br>7<sup>th</sup> April | Country | Deaths in<br>4<sup>th</sup> June |
| --- | --- | --- | --- | --- |
| 1 | Spain | 27.9 | Belgium | 83.1 |
| 2 | Italy | 27.2 | UK | 59.9 |
| 3 | Belgium | 14.6 | Spain | 59.5 |
| 4 | France | 13.6 | Italy | 55.7 |
| 5 | Netherlands | 10.7 | Sweden | 44.4 |
| 6 | UK | 7.9 | France | 43.2 |
| 7 | Luxembourg | 6.8 | Netherlands | 34.6 |
| 8 | Sweden | 4.7 | Ireland | 33.8 |
| 9 | Ireland | 3.5 | Romania | 24.2 |
| 10 | Denmark | 3.2 | Luxembourg | 17.9 |

total country population as of 1<sup>st</sup> of January 2020 (https://ec.europa.eu/eurostat/databrowser/) [19] and COVID-19 total deaths from WHO daily reports [11].

Our search from public health databases showed both Italy and Spain countries rank similarly than other European countries for many of the most common COVID-19 comorbidities. The Italian total population is the world’s second oldest nation while Spain is the 20^th^, with other 16 European countries ahead of it [26], suggesting ageing alone would not explain the similarities observed in these countries. Similarly, the burden of CVD is comparable between Northern Europe (384.3 deaths per 100,000 population) and Spain/Italy (average of 342.5 deaths per 100,000 population [27]). We next check the European diabetes status and today, both type I and type II are higher in North Europe than South Europe [28, 29]. The reported smoking population in Italy and Spain (18.9% and 31.8% respectively) is again similar to other European countries such as Ireland (19.6%), Germany (23.7%) or Portugal (36.8%) [30]. Lastly, data regarding several contributors that lead to liver disease-related deaths from 31 European countries showed the main cause to die from liver disease in Spain and Italy is liver cancer, with the highest hepatocellular carcinoma incidence being observed in the later [31]. Finland’s main cause of liver-disease-related death is alcohol consumption and this could be relevant for the findings explained below. In the same report, it is shown that the prevalence of cirrhosis and other liver disease’s aetiology in Italy is largely due to Hepatitis B and C infections with the highest incidence of hepatitis infections in Europe (5.9%).

### Candidate genes identified and data curation

An extensive literature search was carried out and 291 studies were identified after entering the search terms shown in Figure 1. After removing duplicates, 95 abstracts were screened and a total of 45 publications were included in further pathway analysis due to falling in the inclusion criteria as described previously.

Overall, we identified 63 genes to have a role during a SARS-CoV infection (Supplementary Table 3), with 21 of those being associated with disease aggressiveness and patients’ survival (Supplementary Table 4). In addition, 332 genes have recently been identified in an exclusive study of SARS-CoV-2 − human interactome [32]. These 405 genes were curated as described previously and a total of 437 genes were included in the pathway analysis as a group and as a subgroup, based on the severity of the disease. Empirically and predicted human pathways affected during a SARS infection reported in the literature were identified as well (Supplementary Table 2).

Some of the genes identified, act as ligands or proteases of the viral Spike (S) receptor, assisting in the viral host recognition and entry of the viral genome into the host cell. Genes in this category reported in at least two independent studies have been included in Table 2, while those only identified in one study are in Supplementary Table 1. To mention some, the human transmembrane receptor angiotensin converting enzyme II (ACE2) has been widely reported to be a high-affinity receptor of the SARS-CoV S protein [33–36]. Interestingly, SARS-CoV-2 S protein has shown to have a higher binding affinity to this receptor than SARS-CoV-1 [37]. Several additional human transmembrane receptors and co-activators have also been identified *in vitro* and *in silico* [35, 38–50], indicating SARS-CoV-2 viral RNA entry into the host cell can occur through a different molecular mechanism.

**Table 2:** Host cell candidate genes associated with viral entry.

| Candidate genes | Virus | Host cell response/viral mechanism | Assay methods | Ref. |
| --- | --- | --- | --- | --- |
| **ACE2** | SARS-CoV-1, SARS-CoV-2 | High S binding affinity. Facilitating host cell recognition. | Multiple *in vitro* and *in silico* analysis. | [33-36, 50, 85] |
| **TMPRSS2** | SARS-CoV-1, SARS-CoV-2 | S protein activator, leading to viral membrane conformational change and facilitating SARS virus. | Multiple *in vitro* and *in silico* analysis. | [34, 35, 38, 86] |
| **BSG** | SARS-CoV-2, malaria, HIV, HepB and HHV | *Basigin* genes, which encode CD147 transmembrane glycoprotein recognised by several pathogens. CD147 directly binds to SARS-CoV-2 S protein affecting viral replication. | Review and *in vitro*. SARS-CoV-2 strain isolated from COVID-19 patients. Direct in vitro infection, Co-IP and ELISA. | [41, 42] |
| **HAT** | SARS-CoV-1, HCoV (229E) | *Histone acetyltransferases* family, encoding for a family of cell nuclear enzymes. They contribute to SARS-CoV-1 entry, but it is not essential for S protein activation. | *In vitro*. Both studies: gene cloning, lentiviral expression system, protein expression and cell-cell fusion analysis. | [34, 86] |
| **CLEC4M** | SARS-CoV-1, EVD, Dengue, HCV, CMV, Sindbis, HIV | *C-type lectin domain family 4 member M* genes encode for L-SIGN membrane receptor, recognised by the S protein. Homozygous L-SIGN associated with SARS disease protective role. | *In vitro*. Infection of SARS-CoV-1 human cells, gene expression, cDNA library, IHC assays. Genetic risk association from SARS patients and controls. | [43-46] |
| **ANPEP, ENPEP, DPP4 (or CD26)** | ACE2 studies, HCoV-22944, MERS-CoV45 | Closest co-expression of these three peptidases (R>0.8) with ACE2 in different human tissues. HCoV-22944 binds to ENPEP while MERS to DPP4. | Single cell *in silico* ligand-receptor affinity assays. Data sourced from GEO, Human Cell Atlas, Viral Receptor and Membranome databases. | [38-40] |
| **Cathepsin-B-L** | SARS-CoV-1, SARS-CoV-2, MERS-CoV | Facilitates SARS-CoV-2 cell entry by virus-cell membrane fusion mechanisms but its inhibition does not disable virus entry. | *In vitro*. SARS-CoV-2 S protein pseudovirus system in human lung cell model. | [35, 50] |

ACE2: angiotensin I converting enzyme 2, TMPRSS2: transmembrane Serine Protease 2, ANPEP: alanyl aminopeptidase membrane, ENPEP: glutamyl aminopeptidase, DPP4: dipeptidyl peptidase 4, SARS: Severe acute respiratory syndrome, HIV: human immunodeficiency virus, HepB: hepatitis B virus, HHV: human Herpesviridae, HCoV: human coronavirus, EVD: Ebola virus disease, HCV: hepatitis C virus, CMV: Cytomegalovirus, MERS: middle East respiratory syndrome, S: spike viral protein, CD147 (cluster of differentiation 147), L-SIGN: liver/lymph node-specific intracellular adhesion molecules-3 grabbing non-integrin, Co-IP: co-immunoprecipitation, IHC: immunohistochemistry

Additionally, several candidate human genes were associated with evasion and tuning of the innate and acquired immune system *in vitro* and are summarised on Table 3 and Supplementary Table 1. The genes categorised in this group in at least two independent studies have been included in Table 3, while those only identified in one are on Supplementary Table 1. The main mechanisms of SARS-like virus to escape immune detection and/or processing were by interfering with the hosts’ protein translation [51, 52] (including those associated with antiviral response) and hijacking the immune response [53–64]. They also sequester different host proteins to make precise rearrangements of the endothelium reticulum membrane [34, 63] to build the replication and transcription complex (RTC), effectively creating a protective barrier to replicate, assemble and release viral particles [36, 65, 66] The most common immune signalling pathways affected were those regulated by IFN and NF-κB [55–58].

**Table 3:** Host cell candidate genes associated with viral immune system evasion.

| Candidate genes | Virus | Host cell response/viral mechanism | Research assay | Ref. |
| --- | --- | --- | --- | --- |
| **40s** | Nsp1 studies | Encoding for ribosomal protein S3, interacts with viral Nsp1, inhibiting the host's protein translation by capping the 5'mRNA. | *In vitro*. Reporter gene assays followed by transcriptomics, RNA immunoprecipitation and proteomics assays. | [51, 52] |
| **CCL5, CCL3, CXCL10** | SARS-CoV-1, | These genes encode for IP-10* protein. Increased levels in lung epithelial cells after Nsp1-direct activation of the NF-kB pathway. IP-10 showed specific up-regulation in COVID-19 lung model (when compared to SARS patients). | *In vitro*. Gene cloning, mRNA and protein expression analysis. *Ex vivo*. Lung tissue transfected with COVID-19 and gene expression analysis. | [53, 54] |
| **STING1, TRAF3, TBK1, IKKε** | SARS-CoV-1, HCoV (NL63) | The SARS-CoV-1 PLP transmembrane protein interacts with STING, TRAF3, TBK1 and IKKε disrupting the STING/TBK1/IKKε complex formation and suppressing production of IFN-α and IFN-β, vital for initial innate immune response. PLP protein highly conserved in both SARS-CoV viruses, highlighting the use of potential agonists for this protein as treatments. | *In vitro*. SARS-CoV-1 propagation, and plasmids expressing genes of interest's co-transduction. Co-IP and ubiquitination signalling detection. *In silico*. Homology alignments of both SARS viruses, approved compounds database screening and homology models predictions. | [55, 56] |
| **ADP-ribose** | ssRNA | After binding to Nsp3, post-translational modification of PARP15, PARP14 and PARP10 associated with anti-viral response. | *In vitro*. Cloning, gene expression, mutagenesis, protein purification and crystallization. *In silico*, sequence alignments, glycosylation sites' predictions and 3D mapping. | [62-64] |

40s: 40 subunit, CCL5: C-C motif chemokine ligand 5, CCL3: C-C motif chemokine ligand 3, CXCL10: C-X-C Motif Chemokine Ligand 10, STING1: stimulator of interferon response cGAMP interactor 1, TRAF3: TNF Receptor Associated Factor 3, IKKε: inhibitor of nuclear factor kappa-B kinase subunit epsilon, Nsp1: non-structural protein 1, SARS: Severe acute respiratory syndrome, HCoV: human coronavirus, ssRNA: single strand RNA, \*previously known as IP-10, now CXCL10: C-X-C Motif Chemokine Ligand 10, NF-kB: nuclear factor-kB, PLP: papain-like protein, COVID-19: coronavirus disease 2019, PARP genes: Poly(ADP-Ribose) Polymerase

### Candidate genes associated with pathogenicity

Clinical data from COVID-19 patients’ blood samples have been consolidated in several reviews and retrospective studies, reporting an aberrant chemokine balance in COVID-19 patients (Table 4). While *in vitro* assays suggest this dysregulated production might occur as soon as the S viral protein binds to the host membrane cell receptor [33] there is a correlation between this “cytokine storm” and the severity of the disease [54, 67, 68]. Recent clinical studies report lymphopenia, which it is diagnosed when blood lymphocyte counts are lower than the threshold, to occur early in COVID-19 patients that develop a more severe form of the disease, suggesting its potential application as an early prognostic biomarker [68–72]. Those genes associated with disease aggressiveness too were analysed independently from the rest in the pathway analysis. Additionally, Protein-Protein Interaction map of all the 21 candidate genes associated with COVID-19 aggressiveness, highlighted the physical interactions with lactate dehydrogenase A (LDHA) and LDHB were vital players.

**Table 4:** Candidate genes expressed in SARS and COVID-19 patients and disease severity.

| **Candidate genes** | **Disease** | **Reported observations** | **Ref.** |
| --- | --- | --- | --- |
| ***IL-1B, IFN-γ, CXCL10, and MCP-1*** | SARS and COVID-19 | Increased in SARS and COVID-19 patients. Associated with Th1 cell immunity aberrant response and ARDS | [67] |
| ***IL-4 and IL-10*** | COVID-19 | Increased in COVID-19 patients. Associated with Th2 cell immunity response, facilitating further ARDS | [67] |
| ***IL-2R and IL-6*** | COVID-19 | High expression levels positively correlated with the severity of the disease | [54, 67, 68] |
| ***CXCL10, MCP-1 and TNF-α.*** | COVID-19 | COVID-19 ICU patients have increased serum levels of these genes when compared to non-ICU COVID-19 patients | [54, 71] |
| ***IL-1B, IL-6, cRP*** | COVID-19 | High levels IL-1β of associated with poor prognosis. IL-6 and cRP potential early risk biomarkers | [68] |
| ***Serpine1\**** | COVID-19 | High levels associated with vascular inflammation and higher risk of thrombosis. | [68] |
| ***LDH-hsCRP-lymphocyte*** | COVID-19 | Very accurate (>90%) predictive mortality biomarker signature | [94] |

IL: interleukin, IFN: interferon, MCP-1: monocyte chemoattractant protein 1, CXCL10: C-X-C Motif Chemokine Ligand 10, hsCRP: high-sensitivity C-reactive protein, LDH: lactic dehydrogenase, SARS: Severe acute respiratory syndrome, COVID-19: coronavirus disease 2019, Th: T helper type, ARDS: Acute respiratory distress syndrome, ICU: intensive unit care, \*encodes PAI-I protein

### Signalling pathway analysis

After running the IPA analysis with all the 437 candidate genes, the two most significant canonical signalling pathways were *Caveolar-mediated Endocytosis* (P = 2×10^−19^) and *MSP-RON Signalling* pathways (P = 6.1×10^−19^). *Caveolar-mediated Endocytosis* signalling pathway controls different cellular processes such as endocytosis, cellular signalling and lipid recycling [73]. It regulates the internalisation of different particles, including virus and bacteria, after generating flask-shaped invaginations of the plasma membrane. The *MSP-RON Signalling* pathway contributes to the macrophage induced immune response to assist the host in the viral recognition via the Macrophage Stimulating Protein (MSP) and the transmembrane receptor kinase RON Protein Tyrosine Kinase/Receptor d [74].

On the other hand, when running only those genes associated with COVID-19 survival, the two most significant signalling pathways were *Hepatic Fibrosis and Hepatic Stellate Cell (HSC) Activation* (P = 2.5×10^−18^) and the *Communication between Innate and Adaptive Immune Cells* (P = 1.6.1×10^−14^). The *Hepatic Fibrosis / HSC Activation* occurs in patients with chronic liver damage after accumulating an excess of extracellular matrix (ECM) proteins [75]. This process initiates a series of pro-inflammatory events which downstream effect is the constitutive activation of HSC. When activated, HSCs cells secrete cytokines and their constant activation creates another source of extra ECM, which leaves patients developing tissue fibrosis and liver cirrhosis [75]. Finally, the *Communication between Innate and Adaptive Immune Cells* is the process in which both the immune and adaptive responses interact with each other to defend the host from infection. The initial innate response is crucial as it will determine how strong and specific the adaptive response will be [76].

The two full reports are presented in Supplementary Table 5 and Supplementary Table 6 respectively. The genes associated in these four signalling pathways and their flanking regions can be found in Supplementary Table 7, 8, 9 and 10 and they were further studied in the subsequent PC analysis.

### PC analysis

A total of 81,271,745 variants and 2,504 people passed filters and QC and were therefore included to account and adjust for population stratification for the regions of interest within the genome. The PC analysis results are depicted in Figure 3. Out of four analyses, only the *Hepatic Fibrosis and HSC Activation* (Figure 3c) represented a genetic differentiation for some of the populations tested, with the Spanish and Italian populations clustering closely together and the Finnish population segregating independently in this pathway. The specificity of the Finnish segregation may suggest a protective role of this pathway in this population (5.8 deaths per 100,000 as per 4^th^ of June [11]) or could be a reflection of its well-known unique genetic background [77, 78].

**Figure 3:**
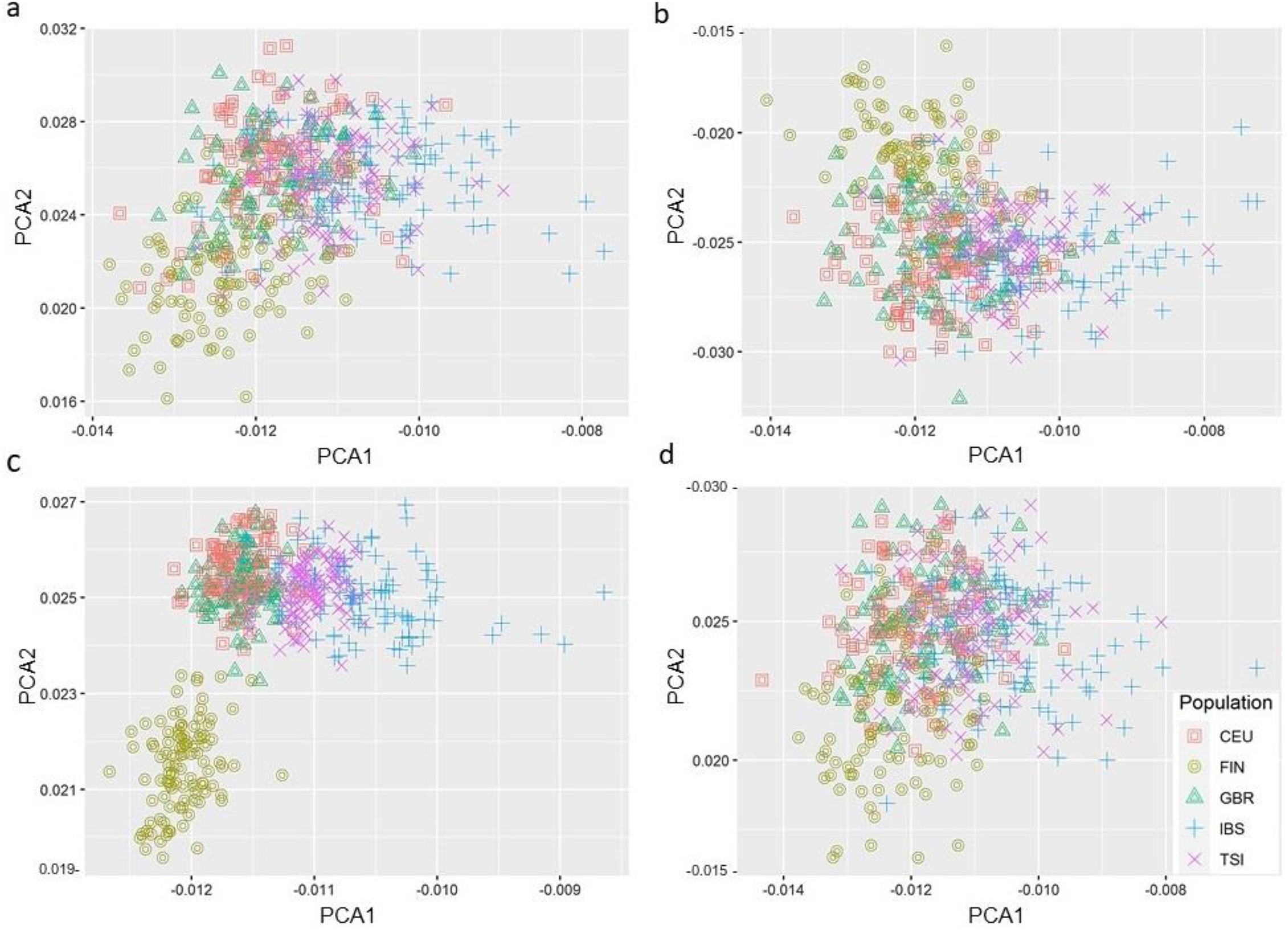
Principal component analysis from five European ancestries for the following canonical signalling pathways: **a**, Caveolar-mediated Endocytosis **b**, MSP-RON Signalling **c**, Hepatic Fibrosis and Hepatic Stellate Cell Activation **d**, Communication between Innate and Adaptive Immune. Population include, CEU: Utah residents with Northern and Western European Ancestry, FIN: Finnish population, GBR: Great Britain (England and Scotland), IBS: Iberian (Iberian Population in Spain), TSI: Tuscany (Tuscany in Italy). Genes obtained from pathway analysis (IPA, Qiagen) and genes locations (GRCh37/hg19 built) included a ±1Mb window. Genome data outsourced from 1,000G phase III v.5b. PC analysis. PC analysis performed using PLINK software v1.9b. Isoforms locations were flanked by ± 1Mb.

## Discussion

We aimed to identify possible genetic variations associated with SARS-CoV viruses’ infections, where allele frequencies are different in the Spanish and Italian populations when compared to other European countries. Despite the strict imposed social restrictions [12] in these two countries, high-mortality rates were maintained for many weeks in the aftermath [11]. Our search from available Public Health data, confirmed the most common COVID-19 comorbidities [13–16, 20, 21] did not fully explain the high mortality rates reported in these two countries, thus we hypothesised that a shared genetic set of traits could partially explain the poor survival outcome. Indeed, genetic studies have shown that, although both populations are closely related to other European populations, they share a higher number of genomic loci [79] while both present certain grades of genomic uniqueness like the diverse haplotypic structure in the Spanish population [80] and the genetic variation inside the Italian population when comparing regions in the country [81]. Also, there are already some well-known genetic variations, in the Italian and Spanish populations associated with disease and disorders. For instance, SNPs in the alcohol dehydrogenase 4 (ADH4) gene are known to play a role in alcohol dependence in the Italian population [82]. Similarly, a Spanish-specific SNP in the ADH1B (rs1229984) and ADH6 (rs3857224) genes are associated with a heavy intake of alcohol amongst this population [83]. In this work, we performed an extensive literature search to identify candidate genes associated with SARS-like viruses’ infection, evasion and hijacking of the immune responses and disease aggressiveness as genetic variations in any of these genes and potential regulatory regions around could have an effect of the pathogeny in vivo. For instance, Chan et al. identified a polymorphism in the gene coding for the extracellular region of the transmembrane receptor L-SIGN (liver/lymph node-specific intracellular adhesion molecules-3 grabbing non-integrin, encoded by *CLEC4M*) that binds to S viral protein to be associated with a lower risk of infections in SARS patients [45]. Similarly, the furin-like cleavage site-specific to the SARS-CoV-2 S protein has been associated with higher pathogenicity *in vitro* [68, 84]. Several host transmembrane receptors have been identified to bind to the S protein [33–36, 41–46, 50, 85] or to prime it [34, 35, 38, 86], with ACE2 and the transmembrane protease serine 2 (TMPRSS2) the two most commonly reported S protein binding receptor and activator respectively. This protease expresses in a larger set of tissues [87] than ACE2 [87–89], suggesting it is the latter which narrows down the infectious cell tropism of the virus. As a note, co-expression studies of ACE2 and TMPRSS2 found it is highest in the nasal epithelium, supporting the observation of a higher viral load detected in nasal than throat swabs in COVID-19 patients and easy droplets viral transmission which could in part explain the high infectious rate [87]. After the S protein is activated, the viral genome enters the host. There are reports of the less well-known membrane trafficking pathway regulator, the two-pore channel subtype 2 (TPC2), which expression is affected by the levels of viral uptake *in vitro* [50], suggesting a role in the pathogenicity *in vivo* similarly to the ssRNA Ebola virus [90]. Our pathway analysis with those genes associated with COVID-19 severity revealed the *Hepatic Fibrosis and HSC Activation* as the most significant canonical pathway. The PC analysis that followed, showed population-specific segregation in the Italian, Spanish and Finnish populations from the rest of the countries analysed in this pathway only. Both the Spanish and Italian populations clustered closely together while the Finnish population segregated independently. These findings, alongside high mortality rate, make this network an area of interest to further investigate treatments for Spanish and Italian COVID-19 patients with current medications for liver disease as an option for those who develop liver malfunction during the infection while those with a pre-existing liver condition may need to go through a more thorough follow up after the recovery period. As mentioned previously, Finland has a unique genetic background originated from geographic and cultural isolation and its population has been the focus of heritable diseases studies and precision medicine treatments [91]. Interestingly, as mentioned above liver cancer is the main cause of liver disease deaths in Spain and Italy while it is alcohol consumption in Finland. This, coupled with the results observed in here, may reflect a correlation between SARS-CoV-2 severe infections in Spanish and Italian patients who suffer not only from liver disease in general but liver cancer in particular while liver damage associated with alcohol consumption does not increase mortality rate as observed in the low mortality rate reported from Finland. It is also possible that the independent opposite segregation of the Finnish population in the *Hepatic Fibrosis / HSC activation signalling* might be a reflection of the unique genetic background previously described. Therefore, we propose that some of the aggressive outcomes observed in COVID-19 patients from Spain and Italy could be partially associated with a pre-existing liver condition and/or may lead to further COVID-19 associated liver complications. Indeed, liver injury in severe COVID-19 patients has been widely reported (review by Zhang C et al. [92]) and SARS-CoV-2 infected patients with symptomatic abnormal liver function has previously accounted for up to 39.4% of the severe cases in a COVID-19 cohort (N = 1,099) [16]. Additionally, although the HSC activation can occur via several molecular and signal pathways [93] its detection could be of special interest in Spain and Italy’s COVID-19 patients. Three biomarkers have recently been identified: lactic dehydrogenase (LDH), lymphocyte and high-sensitivity C-reactive protein (hs-CRP) and when combined they can predict COVID-19 mortality with > 95% accuracy [94]. This could be relevant as LDH can activate HSC [93] and an increase of this biomarker alone (which can predict survival with 92% accuracy on its own) could be an early biomarker of severe cases in these two. On the other hand, there is growing evidence of long non coding RNAs (lncRNAs) associated with SARS-CoV-2 hijacking of mitochondrial DNA and possible lncRNA as key modulators in this [95]. To check this, we asked how many lncRNAs are associated with LDHC which is one of those key proteins regulating liver diseases. We found that NONHSAT15889.1 and NONHSAT018286.2 are the key lncRNAs that are highly expressed in testes from Reads Per Kilobase of the transcripts, per Million mapped reads (RPKM) as available from noncode.org. This agrees with the fact that many men are likely to be near more fatal to death than women, which overlies the liver diseases. As a note, co-expression analysis of ACE2 and TMPRSS2 revealed they co-express in many tissues but not in liver [87], suggesting a different S protein membrane receptors and co-activator mechanism in those severe COVID-19 cases that develop liver dysfunction during the infection. Other efforts to identify several genetic variation in the Spanish and Italian populations associated with severe COVID-19 symptoms have also been reported [96]. However, in this study they focused on potential genes associated with respiratory failure only and these two populations were not analysed in the broader European context.

In terms of future directions, genomic data of other European countries (especially Belgium) and a broader representation of the Italian population should be included in the PC analysis. Validation of the results with a second cohort, preferably with COVID-19 patients, is uncompromising to confirm the results. However, we argue that the recently established *COVID-19 Host Genetics Initiative* will allow us to answer this and many more questions [97].

## Conclusion

In summary, 437 candidate human genes reported to have a role during a SARS virus infection have been identified. Pathway analysis of 21 reported genes associated with severe COVD-19 cases identified *Hepatic Fibrosis and HSC Activation* as the most significant canonical pathway. It was this pathway only where PC analysis showed specific segregation of one or more populations. The Spanish and Italian populations clustered from the rest of the countries while Finnish segregated independently. We propose COVID-19 patients hospitalised in Spain and Italy should be actively surveyed if presenting pre-existing liver conditions, especially those who suffer from liver cancer. Early prognostic biomarkers of liver malfunction in patients with a non-pre-existing condition may play an important role in reducing the death burden observed in these countries.

## Authors Contributions Statement

LM conducted the literature search. LM and PJ searched for candidate genes positions and run pathway analysis. LM prepared the tables and figures. SF carried out PC analysis. PS studied plausible lncRNAs associated with LDHC and conducted protein-protein interaction analysis. LM drafted the manuscript. SF, PS, PJ and JP reviewed and edited the article before submission. JB conceived the initial research hypothesis, provided vital feedback and supervised the work.

## Acknowledgements

We would like to thank you all the team lead by A/Prof Jyotsna Batra for their support during this study and the Cancer Council Australia.

---

**Supplementary table 1:** Candidate genes associated with viral entry and immune evasion reported in one study.

| Candidate genes | Virus | Host cell response/viral mechanism | Research assay | Ref. |
| --- | --- | --- | --- | --- |
| **ADAM17** | SARS-CoV-1 | Competes with TMPRSS2 to cleave ACE2. | *In vitro*. Reporter gene assays, recombinant virus transduction, gene and protein expression analysis. | [34] |
| **CD26 (or DPP4)** | SARS-CoV-2 | Interaction with S1 domain glycoprotein. May play a role in immunoregulation and virulence. | *In silico* structure and glycoprotein binding models. | [47] |
| **DPP4 (or CD26)** | MERS-CoV | After S interaction, it is activated by TMPRSS2 or cathepsin-L. Some isoforms associated with lower viral intake. | *In vitro*. MERS-CoV S cloned in human cell model. Co-IP, flow cytometry and viral titer quantification. | [48] |
| **TNF-α** | SARS-CoV-1 | SARS S protein induces TACE- dependent cleavage of ACE2. Associated with early stage infections. | *In vitro*. Gene cloning and human cell viral infection, gene expression and ACE2 activity assays. | [33] |
| **Furin** | SARS-CoV-2 specific | Furin-like cleave site in the S2 domain associated with increased virulence and viral replication efficiency. | *In vitro*. SARS S protein genomic and proteomics analysis. | [37] |
| **LRRK2, ACSL5, HSD17B4, EPHX1, MCCC2, GSTA4, ACACA, HGD, ROS1, CRIP2** | ACE2 studies | Highest positive expression correlation score with ACE2 in lung ($P \leq 10^{-6}$). *CRIP2*, lowest negative correlation expression ($P = 5 \times 10^{-8}$). | *In silico* assay using RNA-seq lung normal tissue (TCGA), GEO and GO. | [98] |
| **PIKfyve** | SARS-CoV-2 | Main enzyme to synthesize early/late endosome regulators. Assists SARS-CoV-2 pseudovirions *in vitro* entry. | *In vitro*. SARS-CoV-2 S protein pseudovirions expression system in human lung cell model. | [50] |
| **TPC2** | SARS-CoV-2 | Downstream effector of main endosome formation regulator PI(3,5)P2. Assists SARS-CoV-2 pseudovirions *in vitro* entry. | *In vitro*. SARS-CoV-2 S protein pseudovirus expression system in human lung cell model. | [50] |
| **GRP78** | SARS-CoV-2 | Encoded by *Hsp5a* and targeted by PeP42 in cancer cells. S COVID-19 protein model predicts similar binding site. | *In silico* structure and binding predictions of S proteins. Data sourced from HCoV (NCBI), EMBL-EBI. | [49] |
| **CMTR1 (2'-O-MTase)** | SARS-CoV | Predicted to methylate the viral mRNA and, escaping antiviral recognition. | *In silico*. Bioinformatics binding prediction studies. | [99] |
| **IL-8** | SARS-CoV-1 | Increased production after S protein activation. | *In vitro*. Reporter gene assays, recombinant virus transduction in lung epithelial model. EMSA, gene and protein expression assays. | [57] |
| **ubiquitin-aldehyde** | SARS-CoV-2 | Interaction with Nsp3 PLP binding site domain. This highly conserved binding site suggests a vital role in SARS virus. | *In silico*: NCBI Virus repository, MODELLER software, Protein Data Bank, Y2H and literature mining. | [66] |
| **NLRP3** | SARS-CoV-1 | Activated after SARS-CoV-1 E protein calcium ion channel formation. It stimulates inflammasome by overproduction of cytokines. | *In vitro*. SARS-CoV-1 E protein synthetic peptide analysis. | [58] |
| **60s** | Nsp1 studies | After interacting with Nsp1, 60s is suppressed from joining 40s and the 80s translationally competent ribosome cannot translate. | *In vitro*. Expression, translation and rRNA immunoprecipitation and polysome assays. Ribosome binding assay. | [51] |
| **STAT1** | SARS-CoV-1 | SARS-CoV-1 Nsp1 suppresses STAT1 phosphorylation, affecting IFN production. | *In vitro*. Reporter gene assays followed by immunoblot and northern blot analyses. | [59] |
| **PPIA, PPIB, PPIH, PPIG, FKBP1A, FKBP1B** | SARS-CoV-1 | Human cyclophilins/immunophilin that modulate the Calcineurin/NFAT pathway after binding Nsp1. Possibly associated with viral pathogenesis. | *In vitro*. Reporter gene assays followed by SARS-CoV transduction and inhibition assays. | [100] |
| **IRF3** | SARS-CoV-1 | SARS PLP proteases indirectly inhibits IRF3 phosphorylation and its nuclear translocation, suppressing IRF3-dependent IFN production and its pathway. | *In vitro*. Gene cloning, mRNA and protein expression analysis followed by immunoprecipitation. | [60] |
| **TRAF3, IL-1B** | SARS-CoV-1 | After viral ORF3a is translated, it interacts with host's TRAF3, activating the NF-kB pathway and promoting IL-1B transcription. | *In vitro*. Gene cloning, cell transfection, reporter gene and gene expression analysis. | [61] |
| **EDM1, OS9** | SARS-CoV-1 | Nsp3 and Nsp4 interactions initiate RTC synthesis. | *In vitro*. Gene cloning, recombinant BAC-SARS-Nsp3 expression and IP. | [65] |
| **BTS-2** | SARS-CoV-1 | SARS ORF7a inhibits BTS-2 glycosylation, allowing for new virions to be released from host cell. | *In vitro*. Gene cloning, human cells transfection, IP and CD analysis. | [101] |

**Supplementary table 2:**
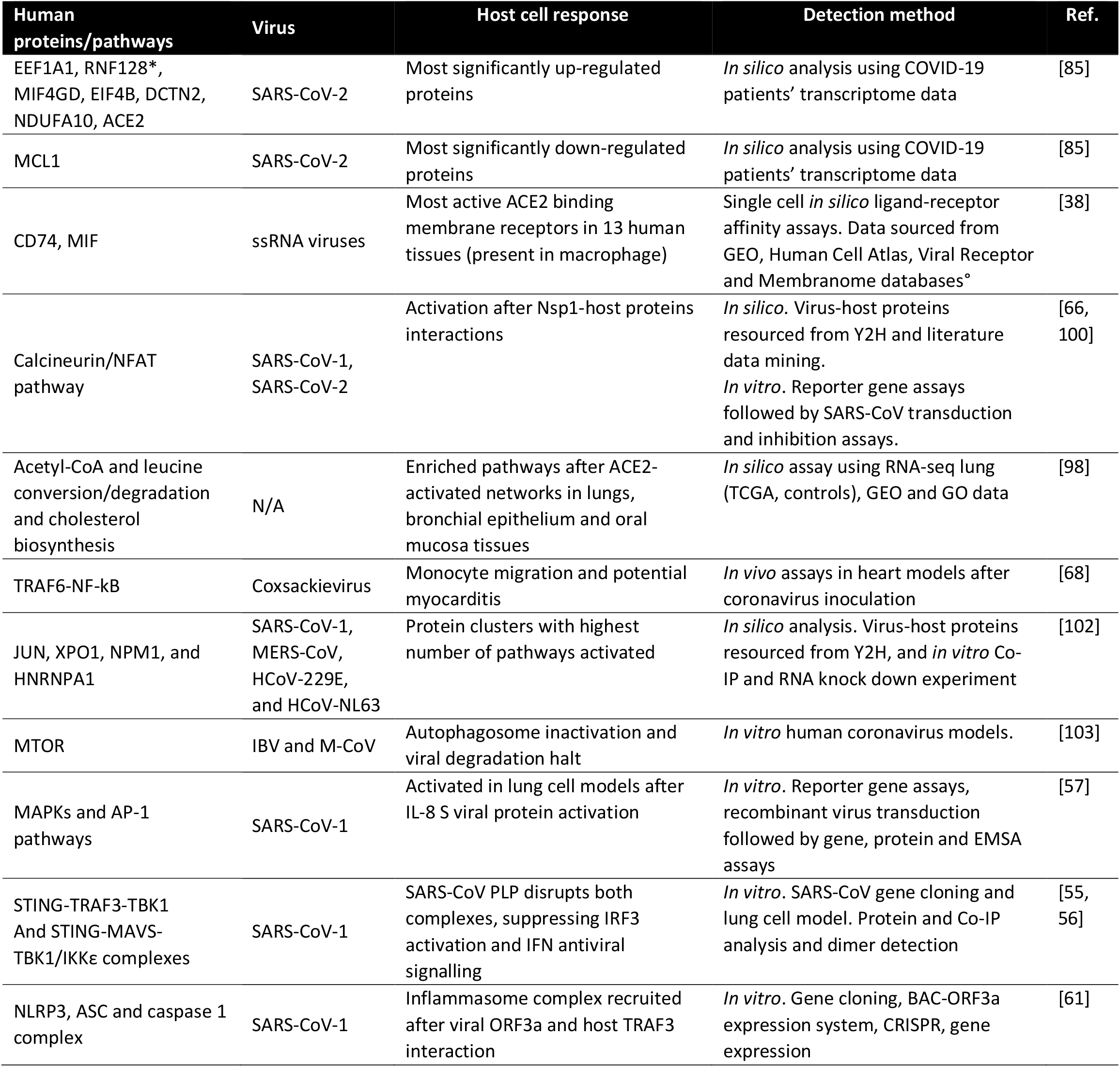
Host cell pathways and protein complexes associated to viral entry and immune system evasion.

| Human proteins/pathways | Virus | Host cell response | Detection method | Ref. |
| --- | --- | --- | --- | --- |
| EEF1A1, RNF128*, MIF4GD, EIF4B, DCTN2, NDUFA10, ACE2 | SARS-CoV-2 | Most significantly up-regulated proteins | *In silico* analysis using COVID-19 patients' transcriptome data | [85] |
| MCL1 | SARS-CoV-2 | Most significantly down-regulated proteins | *In silico* analysis using COVID-19 patients' transcriptome data | [85] |
| CD74, MIF | ssRNA viruses | Most active ACE2 binding membrane receptors in 13 human tissues (present in macrophage) | Single cell *in silico* ligand-receptor affinity assays. Data sourced from GEO, Human Cell Atlas, Viral Receptor and Membranome databases° | [38] |
| Calcineurin/NFAT pathway | SARS-CoV-1, SARS-CoV-2 | Activation after Nsp1-host proteins interactions | *In silico*. Virus-host proteins resourced from Y2H and literature data mining.<br>*In vitro*. Reporter gene assays followed by SARS-CoV transduction and inhibition assays. | [66, 100] |
| Acetyl-CoA and leucine conversion/degradation and cholesterol biosynthesis | N/A | Enriched pathways after ACE2-activated networks in lungs, bronchial epithelium and oral mucosa tissues | *In silico* assay using RNA-seq lung (TCGA, controls), GEO and GO data | [98] |
| TRAF6-NF-kB | Coxsackievirus | Monocyte migration and potential myocarditis | *In vivo* assays in heart models after coronavirus inoculation | [68] |
| JUN, XPO1, NPM1, and HNRNPA1 | SARS-CoV-1, MERS-CoV, HCoV-229E, and HCoV-NL63 | Protein clusters with highest number of pathways activated | *In silico* analysis. Virus-host proteins resourced from Y2H, and *in vitro* Co-IP and RNA knock down experiment | [102] |
| MTOR | IBV and M-CoV | Autophagosome inactivation and viral degradation halt | *In vitro* human coronavirus models. | [103] |
| MAPKs and AP-1 pathways | SARS-CoV-1 | Activated in lung cell models after IL-8 S viral protein activation | *In vitro*. Reporter gene assays, recombinant virus transduction followed by gene, protein and EMSA assays | [57] |
| STING-TRAF3-TBK1 And STING-MAVS-TBK1/IKKε complexes | SARS-CoV-1 | SARS-CoV PLP disrupts both complexes, suppressing IRF3 activation and IFN antiviral signalling | *In vitro*. SARS-CoV gene cloning and lung cell model. Protein and Co-IP analysis and dimer detection | [55, 56] |
| NLRP3, ASC and caspase 1 complex | SARS-CoV-1 | Inflammasome complex recruited after viral ORF3a and host TRAF3 interaction | *In vitro*. Gene cloning, BAC-ORF3a expression system, CRISPR, gene expression | [61] |

