## Supplementary Tables for "Genes associated with liver damage signalling pathways may impact the severity of COVID-19 symptoms in Spanish and Italian populations": Supplementary material.docx

**Supplementary Table 3: Completed list of genes reported to be associated with SARS- viral entry and immune system evasion analysed for pathway analysis.** References presented on Table 2, 3 and Supplementary Table 1.

**Gene symbol**

AAR2

AASS

AATF

ABCC1

ACACA

ACAD9

ACE2

ACSL3

ACSL5

ADAM7

ADAM9

ADAMTS1

AGPS

AKAP8

AKAP8L

AKAP9

ALG11

ALG5

ALG8

ANPEP

AP2A2

AP2M1

ARF6

ARL6IP6

ATE1

ATP13A3

ATP5MG

ATP6AP1

BAG5

BAP1

BCKDK

BCS1L

BSG

BST2

BZW2

C1orf50

CASP1

CCDC86

CCL2

CCL3

CCL5

CD209

CD74

CDH1

CDK5RAP2

CENPF

CEP112

CEP135

CEP250

CEP350

CEP68

CHMP2A

CHPF

CHPF2

CISD3

CIT

CLCC1

CLEC4M

CLIP4

CMTR1

CNTRL

COL6A1

COLGALT1

COMT

CRIP2

CRP

CRTC3

CSDE1

CSNK2A1

CSNK2A2

CSNK2B

CTSL

CUL2

CXCL10

CXCL8

CYB5B

CYB5R3

DCAF7

DCAKD

DCTN2

DCTPP1

DDX10

DDX21

DNAJC11

DNAJC19

DNMT1

DPH5

DPP4

DPY19L1

ECSIT

EDEM1

EDEM3

EEF1A1

EIF4B

EIF4E2

EIF4H

ELOB

ELOC

EMC1

ENPEP

EPHX1

ERC1

ERGIC1

ERLEC1

ERMP1

ERO1B

ERP44

EXOSC2

EXOSC3

EXOSC5

EXOSC8

F2RL1

FAM162A

FAM8A1

FAM98A

FAR2

FBLN5

FBN1

FBN2

FBXL12

FGFR1

FKBP10

FKBP15

FKBP1A

FKBP1B

FKBP7

FOXRED2

FURIN

FYCO1

G3BP1

G3BP2

GCC1

GCC2

GDF15

GFER

GGH

GHITM

GIGYF2

GLA

GNB1

GNG5

GOLGA2

GOLGA3

GOLGA7

GOLGB1

GORASP1

GPAA1

GPX1

GRIPAP1

GRPEL1

GSTA4

GTF2F2

HDAC2

HEATR3

HECTD1

HGD

HMGB1

HMOX1

HNRNPA1

HOOK1

HS2ST1

HS6ST2

HSBP1

HSD17B4

HSPA5

HYOU1

IDE

IFNAR1

IFNAR2

IFNG

IKBKE

IL10

IL17RA

IL1B

IL2RA

IL2RB

IL2RG

IL4

IL6

IMPDH2

INHBE

IRAK4

ITGA1

ITGA10

ITGA11

ITGA2

ITGA2B

ITGA3

ITGA4

ITGA5

ITGA6

ITGA7

ITGA8

ITGA9

ITGB1

ITGB1BP1

ITGB1BP2

ITGB2

ITGB2-AS1

ITGB3

ITGB4

ITGB5

ITGB6

ITGB7

ITGB8

JAKMIP1

JUN

KLK1

KLK10

KLK11

KLK12

KLK13

KLK14

KLK15

KLK2

KLK3

KLK4

KLK5

KLK6

KLK7

KLK8

KLK9

KNG1

LARP1

LARP4B

LARP7

LDHA

LDHB

LDHC

LDHD

LMAN2

LOX

LRRK2

MAP7D1

MARK1

MARK2

MARK3

MAT2B

MAVS

MCCC2

MCL1

MDN1

MEPCE

MFGE8

MIB1

MIF

MIF4GD

MIPOL1

MOGS

MOV10

MPHOSPH10

MRPS2

MRPS25

MRPS27

MRPS5

MTARC1

MTCH1

MYCBP2

NARS2

NAT14

NDFIP2

NDUFA10

NDUFAF1

NDUFAF2

NDUFB9

NEK9

NEU1

NFKB1

NFKB2

NGDN

NGLY1

NIN

NINL

NLRP3

NLRX1

NOL10

NPC2

NPM1

NPTX1

NSD2

NUP210

NUP214

NUP54

NUP58

NUP62

NUP88

NUP98

NUTF2

OS9

PABPC1

PABPC4

PARP10

PARP14

PARP15

PCNT

PCSK6

PDE4DIP

PDZD11

PIGO

PIGS

PIKFYVE

PKP2

PLAT

PLAUR

PLD3

PLEKHA5

PLEKHF2

PLOD2

POFUT1

POGLUT2

POGLUT3

POLA1

POLA2

POR

PPARG

PPIA

PPIB

PPIG

PPIH

PPIL3

PPT1

PRIM1

PRIM2

PRKACA

PRKAR2A

PRKAR2B

PRRC2B

PTBP2

PTGES2

PUSL1

PVR

PYCARD

QSOX2

RAB10

RAB14

RAB18

RAB1A

RAB2A

RAB5C

RAB7A

RAB8A

RAE1

RALA

RAP1GDS1

RBM28

RBM41

RBX1

RDX

RETREG3

RHOA

RIPK1

RNF128

RNF41

ROS1

RPL36

RRP9

RTN4

SBNO1

SCAP

SCARB1

SCCPDH

SDF2

SELENOS

SEPSECS

SERPINE1

SIGMAR1

SIL1

SIRT5

SLC25A21

SLC27A2

SLC30A6

SLC9A3R1

SLU7

SMOC1

SNIP1

SPART

SRP19

SRP54

SRP72

STAT1

STAT2

STC2

STING1

STOML2

SUN2

TAPT1

TARS2

TBCA

TBK1

TBKBP1

TCF12

THTPA

TIMM10

TIMM10B

TIMM29

TIMM8B

TIMM9

TLE1

TLE3

TLE5

TM2D3

TMED5

TMEM39B

TMEM97

TMPRSS11D

TMPRSS2

TNF

TOMM70

TOR1A

TOR1AIP1

TPCN2

TRAF3

TRAF6

TRIM59

TRMT1

TUBGCP2

TYSND1

UBAP2

UBAP2L

UBXN8

UCHL1

UCHL3

UGGT2

UPF1

USP13

USP54

VEGFA

VPS11

VPS39

WASHC4

WFS1

XPO1

ZC3H7A

ZDHHC5

ZNF318

ZNF503

ZYG11B

**Gene symbol**

CRP

IL10

APOA2

IL1B

IL2

CXCL10

IL4

TNF

VEGFA

IL6

SERPINE1

LDHA

LDHC

IFNG

LDHB

CDH1

LDHD

CCL2

PLAUR

IFNAR1

IFNAR2

**Supplementary Table 4: Genes reported to be associated with COVID-19 disease severity and analysed for pathway analysis.** References presented on Table 4.
